# Specificity in transcription factor clustering is encoded in the genome

**DOI:** 10.1101/2024.11.01.621483

**Authors:** Shivali Dongre, Nadine Vastenhouw

**Affiliations:** Center for Integrative Genomics, University of Lausanne, Quartier Sorge, 1015 Lausanne, Switzerland

## Abstract

Transcription factors (TFs) often form clusters in the nucleus. Clusters can facilitate transcription, but it remains unclear how they form. It has been suggested that clusters are seeded by the sequence-specific binding of TFs to DNA, and grow by IDR-IDR interactions that bring in more TFs. This model, however, does not explain how TFs can cluster in specific combinations. Here, we study TF clustering by quantitative imaging of Nanog, Pou5f3, and Sox19b in zebrafish embryos. Using mutant and chimaeric TFs, we show that the formation of a TF cluster requires the DBD as well as at least one of its IDRs. In contrast with the existing model, however, IDRs are not sufficient to join a pre-existing cluster. Instead, both IDR and DBD are needed. Thus, for any TF to join a cluster, motif recognition is required, which explains the specificity in cluster formation. Finally, we show that while IDRs are required to join a cluster, their amino acid sequence is interchangeable, and the DBD can confer specificity to any IDR. Taken together, our work changes the model of cluster formation and explains how specificity is achieved in the organization of transcriptional machinery in the nucleus.

## Introduction

Transcription is a fundamental biological process. To transcribe a gene in the right place and at the right time, several factors need to act in a coordinated manner. Gene-specific TFs recognize and bind to specific DNA sequences within promoter or enhancer regions, and then recruit transcriptional co-activators and general TFs to activate transcription. Gene-specific TFs, cofactors, general TFs, and RNA Polymerase II (RNAPII) are often concentrated in the nucleus in what have been called hubs, clusters, condensates, or transcription bodies (1–9). These clusters are often found at regulatory elements like enhancers (10–12). It has been proposed that the local increase in protein concentration in clusters accelerates biochemical reactions (13) and in line with this, clustering of transcriptional machinery at regulatory elements has been correlated with transcriptional activity ((14–18).

Gene-specific TFs are generally composed of distinct functional units, such as a DNA-binding domain (DBD, *e.g.* homeodomain, zinc finger domain) and a transactivation domain (TAD). The DBD is generally well-structured, allowing for specific recognition and binding to target DNA sequences (19). Transactivation domains, in contrast, often lack a fixed 3D conformation and are part of the intrinsically disordered regions (IDRs) of TFs (20, 21). The disordered nature of TADs allows TFs to interact with a variety of proteins like co-activators, chromatin remodelers, and components of the general transcription machinery. IDRs often extend beyond the TAD and can serve various functions beyond transcriptional activation, such as molecular signaling or serving as scaffolds for protein complexes (19). The structural flexibility of IDRs is crucial for the dynamic and versatile nature of transcriptional regulation, as it enables TFs to form transient and multivalent interactions.

A prevalent model for the clustering of transcriptional machinery postulates that TFs bind to DNA with their DBD and recruit additional molecules into the cluster by IDR-IDR interactions (11, 12, 22, 23). If IDR-IDR interactions are non-specific, this model would predict that all TFs (and other nuclear proteins with an IDR) would cluster together. This is not the case, as different TFs form distinct clusters and co-cluster in distinct patterns (11, 14, 24–28). This calls into question the current model and raises the question how TFs cluster.

Here we used zebrafish embryos to investigate TF clustering. The three gene-specific TFs Nanog, Pou5f3 and Sox19b are required to activate transcription in the zebrafish embryo, and we and others have previously shown that they form clusters in the nucleus (27, 29–32). The availability of mutant embryos that lack these TFs, as well as the ease of supplementing these with synthetic mRNAs coding for mutant forms of the same TFs, make the zebrafish embryo an excellent model to study how the different domains of TFs contribute to cluster formation. Using quantitative live imaging, we find that Nanog-, Pou5f3- and Sox19b-clusters in the nucleus are DNA-seeded. The number of clusters scales with the amount of TF and high affinity sites form clusters first. Our structure-function analysis revealed that the DBD and at least one IDR are required to form a cluster, which is in line with the existing model. Remarkably, however, we find that to join a pre-existing cluster, IDRs are not sufficient and the DBD is also required. Finally, using chimaeric proteins, we show that while IDRs are required to join a cluster, their amino acid sequence is interchangeable, and the DBD can confer specificity to any IDR. Taken together, our results suggest a model where the specificity of TF clustering is determined by the genome.

## Results

To investigate the clustering of TFs, we focused on Nanog, Pou5f3 and Sox19b, all of which play important roles in the activation of transcription in the zebrafish embryo (29–32). We injected RNA encoding each of these TFs fused to mNeonGreen (mNG) into 1-cell stage WT embryos (Figure 1A). For Nanog and Pou5f3, we used the concentration that was shown to rescue the maternal zygotic (MZ) mutant phenotype (30, 36). For Sox19b, the MZ mutant is viable, so here we used the same number of moles of RNA as used for Nanog, for consistency. We visualised the distribution of TFs in the nucleus by spinning disk microscopy (Figure 1B, Diploid). All three TFs form clusters in the nucleus as previously reported (18, 27, 42, 43). Quantification of the number of clusters at the 512-cell stage, when clusters are abundant and clearly visible, showed that Nanog, Pou5f3 and Sox19b form distinct numbers of clusters (Figure 1C, diploid embryos). We detect a median of 30 Nanog clusters per nucleus. The number of Pou5f3 clusters is similar with a median of 33, and Sox19b generally forms only two clusters, and only transiently. These numbers lie in a similar range at the 256- and 1k-cell stage (Figure S1).

**Figure 1.**
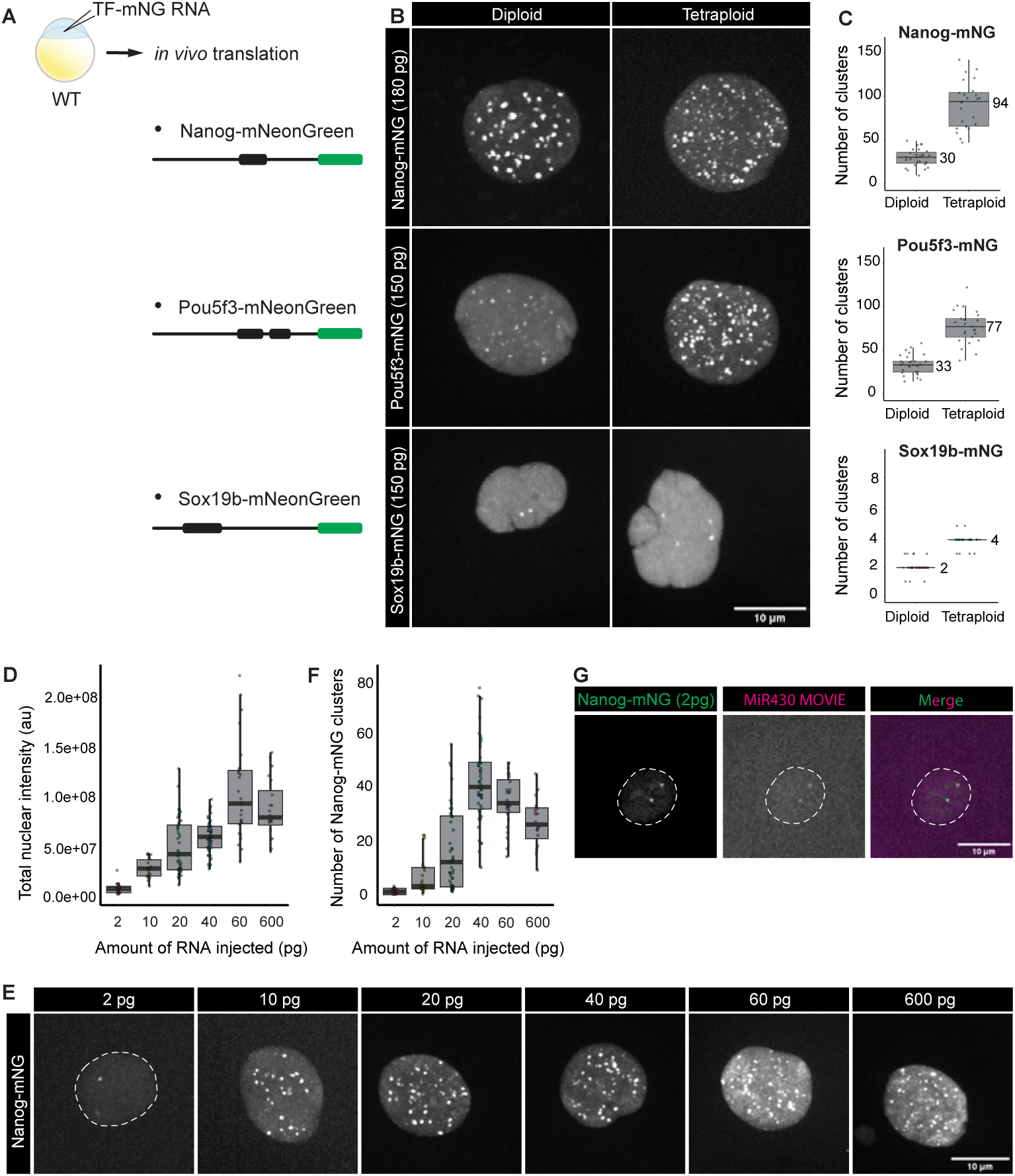
TFs form clusters in the nucleus, and these are DNA-seeded. **A.** RNA coding for Nanog-mNG, Pou5f3-mNG and Sox19b-mNG was injected in 1-cell stage WT embryos. Images were taken at 512-cell stage and at the midpoint between two mitoses, except for Sox19b for which the image was taken right after mitosis because clusters only from transiently. **B.** Visualisation of Nanog-mNG, Sox19B-mNG and Pou5f3-mNG in diploid and tetraploid embryos. **C**. Quantification of the number of clusters per nucleus in diploid and tetraploid WT embryos. The median values of the distributions are indicated in the graphs. Quantifications for additional stages are shown in Figure S1. With N as the number of embryos, and n as the total number of nuclei, N ≥9 and n ≥25 (see Methods for more details on replicates). **D.** Quantification of total nuclear intensity of mNeonGreen in individual nuclei as a proxy for total nuclear Nanog protein at different concentrations of injected RNA. au = arbitrary units. **E.** Visualization of Nanog-mNG clusters after injecting WT embryos with a concentration series of Nanog-mNG RNA, as indicated. **F.** Quantification of the number of clusters per nucleus at different concentrations of injected RNA. With N as the number of embryos and n as the total number of nuclei, N ≥8 and n ≥25. **G.** Visualisation of the overlap between Nanog-mNG and MiR430 RNA in WT embryos that were injected with 2 pg of Nanog-mNG RNA. In B, C and E, Maximum Intensity Projections (MIPs) in Z of representative individual nuclei extracted from spinning disk confocal microscopy at the 512-cell stage are shown.

The concentration of a specific TF in the nucleus has been shown to influence the number of clusters that form (23, 43, 44). To directly test the relationship between TF concentration and cluster number, we injected between 2 and 600 pg of RNA encoding Nanog-mNG into MZ*nanog* embryos. To determine whether the increase in injected RNA resulted in an increase in protein, we then measured the total intensity of nuclear mNeongreen in all conditions, using this as a proxy for the total amount of Nanog protein in the nucleus. While an increase in RNA initially results in an increase in protein, no further increase in the amount of protein is observed between 60 and 600 pg of injected RNA (Figure 1D). We speculate that this might be due to saturation of the translational machinery. We then quantified the number of clusters that formed with an increasing amount of RNA. As we increased from 2 to 40 pg of injected RNA, the number of clusters in the nucleus increases. Above this concentration, the number of clusters stabilized and even decreased (Figure 1E, F). We conclude that an increase in the amount of protein in the nucleus results in an increase in the number of clusters that form.

The clustering of TFs may reflect the formation of protein aggregates in the nucleoplasm (23), or the formation of clusters that are DNA seeded (10, 45–49). To distinguish between these two possibilities, we took advantage of the ease of making tetraploid zebrafish embryos (37, 50) (see Methods). If clusters are TF aggregates, a doubling of DNA would be predicted to have a negligible effect on the number of clusters at the same TF concentration. If clusters are DNA seeded, however, it would be predicted to roughly double the number of TF clusters. We injected tetraploid embryos with the same constructs at the same concentration as the diploid embryos, and observed that the number of clusters increased in tetraploid embryos (Figure 1B). Quantification of clusters showed approximately a doubling of the number of clusters for Sox19b and Pou5f3 (Figure 1C), while the number of Nanog clusters nearly tripled. These results show that the number of clusters is correlated with the DNA content. We conclude that the Nanog, Pou5f3 and Sox19b clusters are DNA-seeded.

Remarkably, we see two Nanog clusters even when injecting only 2 pg of RNA (Figure 1E). We and others have previously shown that two of the Nanog clusters in the nucleus form at the *mir430* locus (18, 27, 42). This locus has many binding sites for Nanog (29, 51) and can thus be considered a high affinity region. In line with this, the Nanog clusters that form on the *mir430* locus are the first to appear when cells exit mitosis(18). We therefore hypothesized that the two Nanog clusters that form when injecting 2 pg of Nanog RNA form at the *mir430* locus. Visualisation of MiR430 transcripts using Morpholino VIsualization of Expression (34) (MoVIE) confirmed this (Figure 1G). We conclude that the concentration of a specific TF affects the number of clusters that form, and that factors cluster first at high affinity sites.

The model for TF clustering that has been put forward postulates that TFs bind sequence-specifically to DNA, after which additional factors can join by means of their IDR. In the very simple scenario in which there is no specificity in the interactions between IDRs, this would predict that once a TF is bound to DNA, any other protein with an IDR could associate with this TF and contribute to cluster formation. If this were the case, TF clusters would be predicted to contain all TFs with IDRs. To directly assess the specificity in TF clustering, we asked whether the Nanog, Sox19B and Pou5f3 clusters always overlap (non-specific clustering) or form clusters in different combinations (specific clustering). To do so, we generated the constructs shown in Figure 1A but now with the fluorescent protein mScarlet (mSc), which does not spectrally overlap with mNeongreen. Co-injecting the mNeongreen and mScarlet constructs, we could visualise two TFs at the same time and assess their co-localization (Figure 2). Sox19b-mSc forms two clusters (as also seen with mNG, Figure 1B) and these colocalize with Nanog as well as with Pou5f3 (empty arrowheads). These are the clusters that form at the *mir430* locus (27). Both Nanog and Pou5f3, however, form many more clusters, and these do not colocalize with Sox19b (filled arrowheads). Similarly, when assessing the overlap between Nanog and Pou5f3 clusters, we observe clusters that are Nanog-only and Pou5f3-only (Figure 2A (filled arrowheads), 2B). We conclude that there is a high level of specificity in the co-clustering of Nanog, Sox19b and Pou5f3. This is an agreement with the specificity in TF cluster formation that others have observed (6, 52, 53).

**Figure 2.**
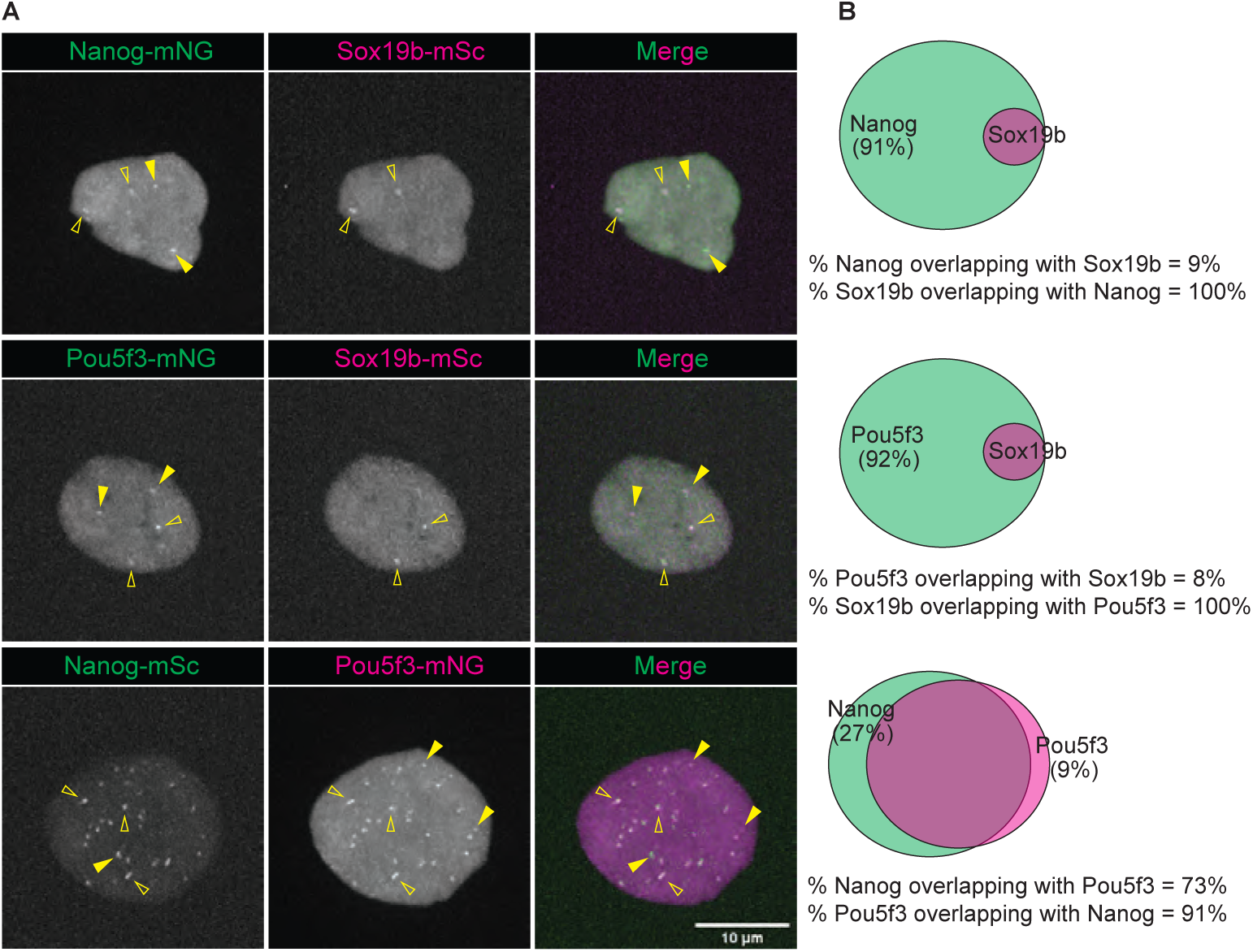
Nanog, Sox19b and Pou5f3 cluster in specific combinations. **A.** Pairwise visualisation of TFs as indicated. Shown are representative images of Maximum Intensity Projections (MIPs) in z of individual nuclei extracted from spinning disk confocal microscopy at 512-cell stage. Images were taken at the midpoint between two mitoses, except in comparisons including Sox19b for which the image were taken right after mitosis because Sox19b clusters only from transiently. Non-overlapping examples are indicated with filled arrowheads and overlapping examples are indicated with empty arrowheads. **B.** Venn diagrams depict the median values of the fraction of overlapping clusters for pairwise co-labelled TFs. Quantification of overlap was done at the midpoint between two mitoses, when all the clusters have formed and are clearly visible, except in cases involving Sox19b (see Methods for details). The percentages in the Venn diagrams represent the non-overlapping fraction of clusters. With N as the number of embryos and n as the number of nuclei, N ≥8 and n ≥25.

We then set out to investigate what mediates the specificity in clustering of Nanog, Sox19b and Pou5f3. The three proteins show the typical structure of a TF with a highly structured DBD and disordered regions of variable length that flank this domain (Figure S2A). The Nanog DBD (homeodomain (HD)) is flanked by two IDRs of similar size. The Sox19b DBD (HMG-domain) is also flanked by two IDRs: one shorter N-terminal IDR, and a longer C-terminal IDR. Similarly, the Pou5f3 DBD (there are two: Pou and homeodomain), is flanked by a long N-terminal IDR and a shorter C-terminal IDR. To investigate the role of DBD and IDRs in cluster formation, we created constructs in which each of them was deleted or mutated (Figure 3A, Figure S2B).

**Figure 3.**
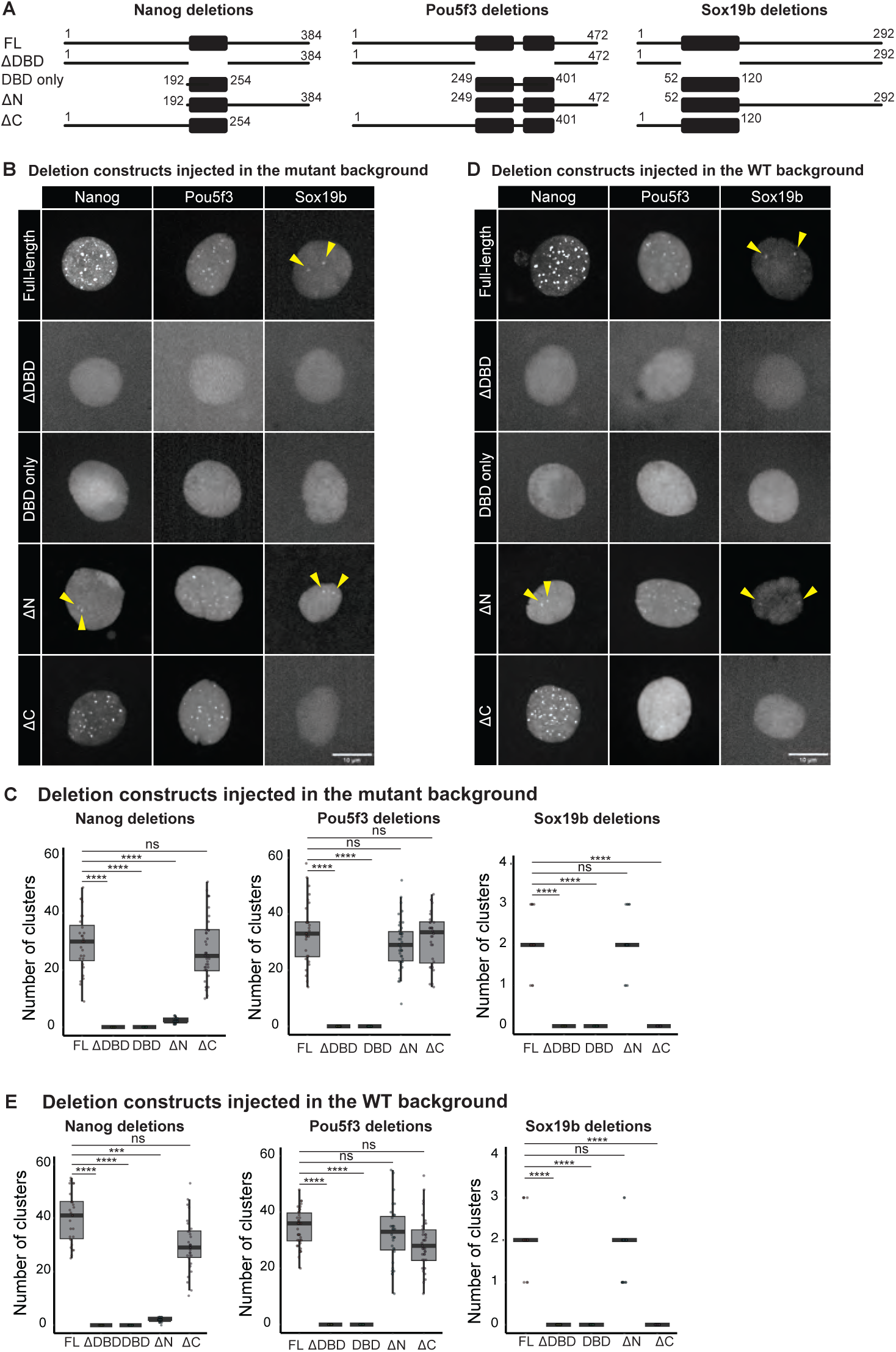
DBD is required to join an existing cluster. **A.** Schematic representation of the deletion constructs that were injected into 1-cell stage embryos. Numbers indicate the amino acids that were retained. DBD is shown as a black box. **B.** Images obtained after injection of the indicated constructs in the respective TF MZ mutants. Yellow arrowheads indicate the two clusters formed by NanogΔN-mNG, full-length Sox19b and Sox19b deletions. Shown are representative images of MIPs of individual nuclei at interphase extracted from spinning disk confocal microscopy at 512-cell stage. Images were taken at the midpoint between two mitoses, except for full-length Sox19b, Sox19b deletions and NanogΔN-mNG, for which the image was taken right after mitosis because these clusters only from transiently. **C.** Quantification of the number of clusters shown in panel B. With N as the number of embryos and n as the number of nuclei, N ≥7 and n ≥28. **D.** As in D, but after injection of the indicated constructs injected in WT embryos. **E.** Quantification of the number of clusters represented in panel D. With N as the number of embryos and n as the number of total nuclei, N ≥7 and n ≥28. Statistical analysis in C and E was performed using Kruskal Wallis test with Dunn’s multiple comparisons.

We distinguish the seeding of a cluster (which refers to the binding of a TF to the DNA) from the growth of a cluster (which refers to the joining of additional factors to an already existing cluster). First, to investigate which domains are important to form a cluster (both seeding and growth), we injected the constructs we generated in embryos that are MZ mutant for the injected TF so that no endogenous full-length protein is present to facilitate clustering (30, 36). We first injected Nanog without its DNA binding domain (ΔDBD) in MZ*nanog* embryos. This did not result in the formation of any clusters (Figure 3B,C). This shows that, as expected, the DBD is required for the formation of Nanog clusters. This is specifically due to the loss of its DNA-binding capacity, because a point-mutant that abrogates DNA-binding gives the same result (Figure S2B,C). The DBD, while required, and by itself sufficient to bind to DNA as evidenced by its mitotic retention (Figure S3), is not sufficient for cluster formation, because we see no clusters when we inject the Nanog DBD-only construct (Figure 3B,C). Thus, both the DBD as well as the IDRs are required for cluster formation. This is in line with the existing model. We then tested the role of the N- and C-terminal IDR of Nanog separately. Interestingly, only two clusters per nucleus were observed on average (Figure 3B,C). These form at the *mir430* locus (Figure S2D). Because the N-terminal IDR has been shown to harbour a dimerization domain (54), however, it is unclear whether the importance of this IDR is caused by the loss of a single IDR or the loss of the dimerization domain. We therefore turned to the results obtained after deletion of the C-terminal IDR. Here, we find that clusters still form and their numbers are comparable with those for full-length Nanog (Figure 3B,C). We therefore conclude that for Nanog, an IDR or dimerization domain in addition to the DBD, is sufficient to form clusters. We then injected different variants of Pou5f3 in MZ*pou5f3* mutant embryos (36). Again, we find that DBD as well as IDRs are important for clustering (Figure 3B,C). Here, both IDRs are individually dispensable for clustering, suggesting that one IDR in addition to the DBD, is sufficient to generate clusters. Finally, we repeated these experiments for Sox19b, using MZ*sox19b* embryos (36). These experiments confirmed that both the DBD as well as IDRs are important for clustering. Interestingly, in this case, the deletion of the short IDR (N-terminal) had no effect on clustering, while the removal of the longer IDR (C-terminal) abrogated clustering (Figure 3B,C). This might suggest that IDR length matters for the ability of TFs to cluster. We note here though, that even full-length Sox19b forms only two clusters, so the dynamic range of our analysis is not as good as for Nanog and Pou5f3. Taken together, we conclude that both DBD and at least one IDR are required to form a cluster. This is in line with a model in which the DBD is required to seed a cluster, and IDRs are required to grow it.

Next, we set out to investigate whether IDRs are sufficient to join a cluster, as was suggested before (11, 23). If true, it would be expected that any TF that retains either one or both IDRs would integrate into an existing cluster. To test this, we repeated the experiment described above, but now instead of injecting the constructs into mutant embryos, we used WT embryos, in which clusters of endogenous full-length protein are present (42, 43). As expected, full-length Nanog is seen in clusters (Figure 3D,E). The number of clusters lies in the same range as was seen for the mutant embryos (Figure 3C, E), suggesting that the that the injected Nanog joins existing clusters. The homeodomain only (lacking both IDRs), in contrast, was unable to join clusters. The presence of an IDR along with the homeodomain restored the ability to join a cluster, with - similar to above - a stronger effect of the N- than the C-terminal IDR (Figure 3D,E). Thus, to join a cluster, IDRs are indispensable, as would be predicted by the model.

Contrary to expectations, however, we observed that neither NanogΔHD nor Nanog HD R247A were able to join existing clusters (Figure 3D,E and Figure S2E). This shows that the Nanog DBD is required for Nanog to join an existing cluster and IDR-IDR interactions alone are not sufficient. Experiments with Sox19b and Pou5f3 confirmed this (Figure 3D,E). We conclude that the DBD of TFs is required even to join an existing cluster, which is in contrast with the current model for TF clustering.

The observation that the DBD of a TF is required to join an existing TF cluster suggests that specificity in cluster formation is encoded by the DBD. To test this directly, we generated chimaeric TFs in which the Nanog DBD is flanked by the IDRs of Sox19b (referred to as SNS), or the RNA binding protein Fus (referred to as FNF) (Figure 4A). We chose the Sox19b IDRs because we have shown that it co-clusters with Nanog (Figure 2), indicating that the IDRs of these two TFs are compatible. We selected the Fus IDRs because they have been shown to facilitate clustering when fused to proteins (16), yet full-length zebrafish Fus does not form any clusters in the nucleus at 512-cell stage (Figure S4). These chimaeric TFs were fused to mNeonGreen as before (Figure 4A). We injected the chimaeras and compared their clustering in MZ*nanog* embryos with that of full-length Nanog (Figure 4B, C). We observed that the clustering of the SNS and FNF chimaeras show a stark difference from the clustering pattern of full-length Sox19b and full-length Fus (Figure 1B, S4). Rather, the chimaeric proteins cluster like full-length Nanog (Figure 4B), as confirmed by the quantification of the number of clusters (Figure 4C). To further test this, we generated a chimaera that contained a Sox19bDBD flanked by Nanog IDRs (NSN). This chimaera formed only two clusters, which is the same as we observed for full-length Sox19b (Figure S5). Together this confirms that specificity in clustering is mediated by the DNA binding domain, and shows that IDRs play a sequence non-specific role in clustering.

**Figure 4.**
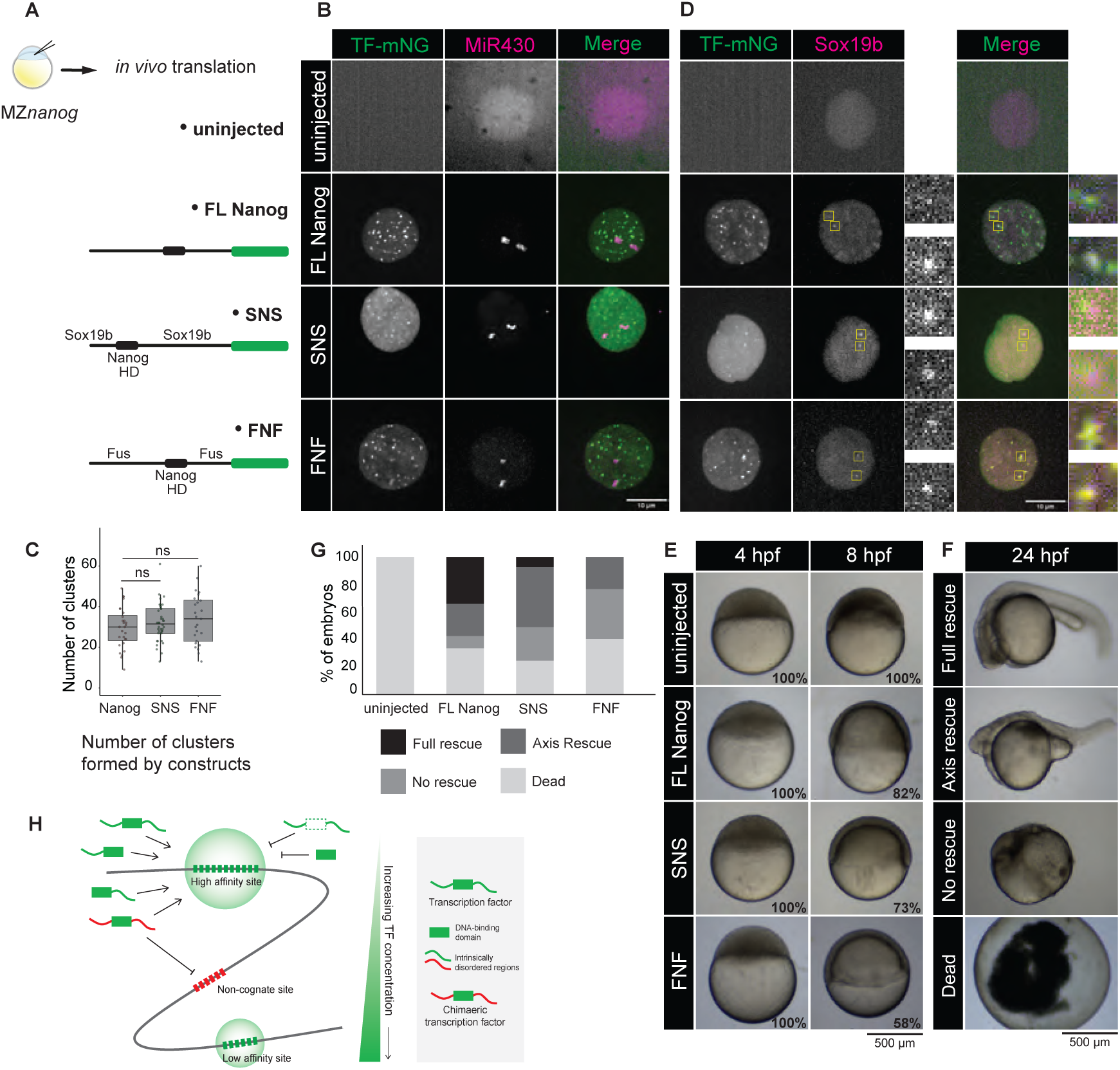
Specificity in TF clustering is mediated by the DBD. **A.** RNA coding for indicated protein was injected in 1-cell stage MZnanog embryos. **B.** Visualisation of the indicated TFs and MiR430 RNA in MZnanog embryos. Shown are representative images of MIPs of individual nuclei at interphase extracted from spinning disk confocal microscopy at 512-cell stage. **C.** Quantification of the number of Nanog-mNG, SNS-mNG, or FNF-mNG clusters per nucleus in MZnanog embryos. With N as the number of embryos, and n as the number of total nuclei where N ≥9 and n ≥25. Statistical analysis was performed using Kruskal Willis test with Dunn’s multiple comparisons. **D**. Visualisation of the indicated TFs and Sox19b in MZnanog embryos. The yellow boxed regions highlight the two Sox19b clusters, with the inset providing a zoomed-in view. Shown are representative images of MIPs as in B. **E**. Embryonic phenotypes of MZnanog embryos injected with the indicated RNA. Shown are representative images of individual embryos captured at 4 hpf and 8 hpf. The percentages indicate the fraction of embryos with the shown phenotype, which is also observed in time matched WT embryos. **F.** Classification of the embryonic phenotypes observed at 24 hpf, as described in Veil et al., 2018 (30). **G.** Classification of embryos at 24hpf upon injection of chimaeric constructs. Categories are described in F. No uninjected embryos survived to reach 24 hpf, they all died during epiboly. With N as the number of embryos, and n as the number of total embryos, N=3 and n>180 embryos. **H.** Schematic model for TF clustering. TFs cluster in a concentration dependent manner, forming first at high affinity sites. Both DBD and IDR are required to form a cluster. The DBD is dictates where clusters form. IDRs are necessary for clustering, but by themselves, cannot join a cluster.

In the chimaera experiments, two of the SNS and FNF clusters form at the *mir430* locus, as visualised by labelling MiR430 transcripts (Figure 4B). We and others have previously shown that Nanog is important to activate transcription from the *mir430* locus (27, 29). The fact that we can see MiR430 transcripts when we inject chimaeric proteins in a MZ*nanog* background shows that these chimaeric proteins are not only able to cluster, but also to activate transcription of the *mir430* locus, which makes them functionally similar to full-length Nanog. We have previously shown that at this locus, Nanog recruits Sox19b to activate transcription (27). This would suggest that the chimaeric proteins SNS and FNF are also able to recruit Sox19b. This is indeed the case because the SNS and FNF clusters both colocalize with Sox19b clusters (Figure 4D, yellow boxes). Finally, we tested whether the chimaeric constructs can also rescue the developmental arrest early during gastrulation seen in MZ*nanog* mutants. Interestingly, both the SNS and FNF chimaeric constructs rescue the gastrulation phenotype, suggesting that they can functionally replace full-length Nanog not only at the *mir430* locus, but also at other Nanog target genes (Figure 4E). We have previously found that neither the Nanog-DBD-only nor the Nanog **Δ**DBD can rescue MZ*nanog* phenotype (27), so while the specific sequence of the IDRs does not seem to matter, the presence of the Nanog DBD and at least one IDR is required for the functional rescue at this developmental stage. The chimaeras, however, fail to completely rescue the mutant phenotype at later stages of development (Figure 4F,G).

This may suggest that later in development the sequence of the IDR does matter, but we cannot exclude other possibilities such as, for example, the faster degradation of SNS and FNF than FL Nanog. We conclude that the IDR has a non-sequence specific role in clustering in early embryogenesis.

## Discussion

In this study, we show that the TFs Nanog, Pou5f3 and Sox19b form clusters in the nucleus, that these clusters are DNA seeded, and that their number scales with the amount of TF. As previously proposed, the DBD as well as at least one IDR are necessary to form a cluster. Surprisingly, however, the DBD is also required to join an existing cluster. Using chimaeras, we confirm that the DBD determines the specificity of cluster formation and show that IDRs contribute to clustering through multivalent interactions, independent of their amino acid sequence.

### Binding site affinity and IDR-IDR interactions together promote the crowding needed for clustering

Our data shows that both the DBD and the IDR of a TF are required for the formation of clusters on DNA (Figure 3B, C). A widely used model for TF clustering postulates that TFs bind to DNA via their DNA binding domain, followed by the recruitment of additional factors, facilitated by IDR-IDR interactions between proteins (3, 11, 22, 55, 56). The importance of the DBD in clustering is undisputed when focusing on DNA-bound clusters (10, 45–48), although it might be dispensable when protein-protein interactions are abundant, as is - for example - the case for Zelda when it is recruited to GAF-rich regions in the *Drosophila* embryo (57). The role of IDRs in the formation of DNA-bound TF clusters is less well understood. While IDRs certainly facilitate the clustering of TFs both *in vitro* (11, 12, 58) and *in vivo* (4, 16), they are not always required for cluster formation (45, 46, 49, 53, 59). When IDRs are dispensable, the clusters are often seeded by a high density of closely spaced binding sites (45, 49, 59), which can probably bring together many proteins without the need of IDR-mediated interactions. Indeed, the number of TF binding sites has been shown to facilitate TF cluster formation (22, 60, 61). Related to this, our work shows that the number of TF clusters that form depends on the concentration of a TF (Figure 1E, F). This seems intuitive, and indeed, previous work in *S. cerevisiae* had found a similar relationship between protein concentration and the clustering of the yeast TF Gal4 (23). In this study, however, not all the Gal4 clusters were associated with DNA, which made it difficult to uncouple the behaviour of DNA seeded clusters from non-DNA bound clusters, especially because *in vitro* work has shown that the formation of non DNA-bound clusters is also concentration dependent (22, 53). In our study, Nanog clusters are DNA-bound (Figure 1B,C, Figure 3B,C, Figure S2C). These DNA-bound TF clusters form preferentially at high affinity binding sites (Figure 1G), and, with increasing TF concentration, also form at lower affinity sites. Taken together, this shows that the crowding that is needed for TF clustering is promoted by high numbers of binding sites as well as protein-protein interactions, mediated by IDRs.

### Specificity in cluster formation is encoded in the genome

Our data shows that once a TF cluster has been seeded, additional TFs can join the cluster only when a functional DBD is present (Figure 3 D, E). This is in contrast with current models which propose that TFs first bind to DNA via their DBD, followed by IDR-mediated recruitment of additional factors to create a cluster (3, 11, 22, 23) as this model would predict that once a cluster has been seeded, additional molecules can join the cluster through IDR-IDR interactions. Our data shows that this is not the case for gene-specific TFs (Figure 3). Thus, even though not all TF in a cluster are bound to DNA at any given time (19, 27, 61), they all need to possess a functional DBD to join the cluster (Figure 4H). This result implies that when co-clustering of different TFs is observed (6, 52, 53), this is driven by the presence of binding sites for the factors that are seen to co-cluster, and not simply by IDR-IDR interactions. Indeed, at the *mir430* locus, where we see co-clustering of Nanog, Sox19b and Pou5f3, binding sites for these factors have been identified (29, 34). Extrapolating this to other clusters where multiple TFs come together, we assume that these represent other genomic regions where binding sites for these factors co-occur. We conclude that the clustering of TFs is dictated by the genome, which ensures specificity in TF cluster formation (Figure 4H).

### The role of IDRs is different for the clustering of gene-specific and general TFs

Our chimaera experiments show that IDRs of gene-specific TFs facilitate clustering, but that their specific protein sequence is not important for clustering (Figure 4). We note here that the Nanog and Sox19b IDRs that we tested in our chimaeras, are in principle compatible, as they can co-cluster endogenously. Fus, however, does not form clusters in nuclei of zebrafish embryos at the stage we analyzed. Therefore, especially the ability of Fus IDRs to functionally replace Nanog IDRs suggests that IDRs have a sequence non-specific role in cluster formation and transcription activation in the early embryo. It has been shown that IDRs can mediate specificity in the binding of TF to DNA (62–64), so we cannot exclude the possibility that our observation is limited to early embryonic stages. This may be supported by our observation that the chimaeric TFs can recapitulate the function of the WT TFs in the early embryo, but fail to do so at later stages of development (Figure 4 E-G). Our results are in contrast with data on the clustering of general TFs (58, 65). These studies show that IDR-IDR interactions can be selective (58) and that this specificity in co-clustering can be mediated by patterns of charged residues, arranged in alternating blocks within the IDRs (65). Thus, while IDRs do not seem to provide specificity when gene-specific TFs cluster, they do provide specificity in the clustering of general TFs. We conclude that gene-specific TFs cluster specifically at sites that are recognized by their DBD. IDRs are important to facilitate clustering but are not sufficient for a gene-specific TF to join an existing cluster. To form a competent transcription body, general TFs such as Mediator and RNA pol II are recruited to these clusters. For these factors, IDRs seem to be sufficient to join a cluster. This would ensure the sequence-specific clustering of gene-specific TFs, while facilitating the recruitment of general TFs that are universally required to initiate transcription.

## Materials and Methods

### Zebrafish husbandry and manipulation

Zebrafish were maintained and raised under standard conditions, and according to Swiss regulations (canton Vaud, licence number VD-H28). Wild type (TLAB) and mutant (*nanog^−/−^*, *sox19b^−/−^*, *pou5f3^−/−^*) fish were used for this study. Embryos were collected immediately after fertilisation. The embryos were grown at 28℃ and the developmental stage was determined as described by Kimmel (33). Developmental stages of mutant and injected embryos were determined by comparing them to wild type embryos obtained at the same time and grown under identical conditions.

### Microinjection of RNA and morpholinos

To visualize TFs (full-length, mutated, and chimaeric), embryos were injected at the 1-cell stage with the mRNA coding for a specific TF-fluorophore fusion. The fusion mRNA was translated within the embryo and the resulting protein visualized by spinning disc microscopy. The *in vitro* synthesized mRNA for the TF-fluorophore fusion constructs were injected at the concentration and molar amount that is indicated in the table below. We ensured that the molar amount of Nanog, Pou5f3 and Sox19b deletion constructs as well as the SNS, FNF and NSN chimaeras was equivalent to the full-length construct containing the corresponding DBD. To label nascent MiR430 RNA, Lissamine-labeled anti-MiR430 morpholino (5’-TCTACCCCAACTTGATAGCACTTTC-3’-Lissamine, Gene Tools) was injected in 1-cell stage embryos at 14 fmole/embryo as described previously (34).

### Amount of RNA injected for each of the constructs

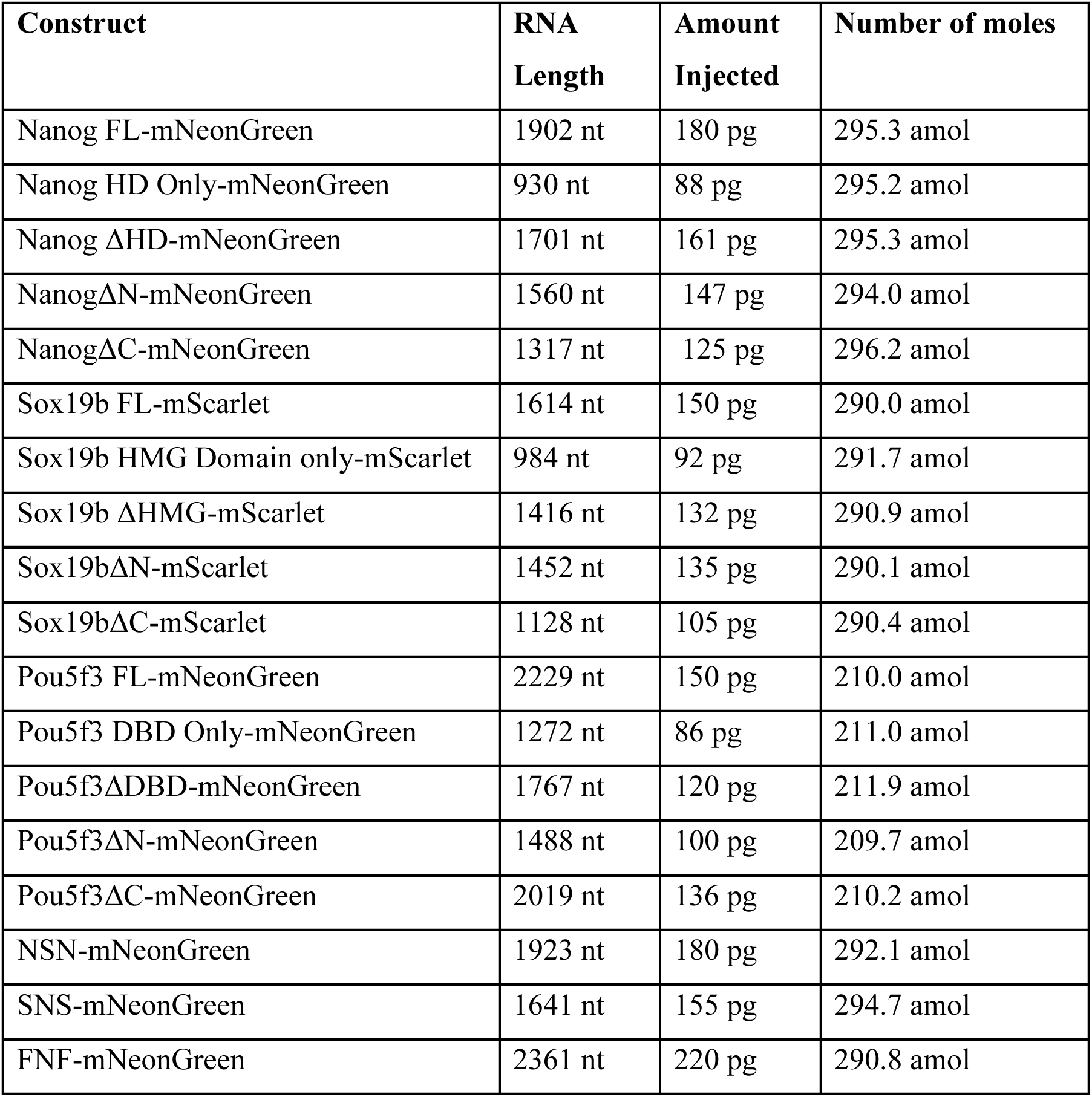

### Generation of deletion constructs and chimaeras

Full-length plasmids for Nanog, Pou5f3 and Sox19b in the pCS2+ vector, tagged with mNeonGreen or mScarlet at the C-terminus, were obtained from prior work (27). Deletion constructs for Nanog were previously established (27), while those for Pou5f3 and Sox19b were generated using the plasmids with the sequence encoding the full-lengh proteins as templates. All around PCR was performed on the full-length constructs in pCS2+, by using primers flanking the segment which were to be deleted. The exact number of amino acids that are deleted are mentioned in Figure 3A. To make the chimaeras, the individual segments were amplified by PCR and cloned using the Gibson Assembly kit and following the manufacturer’s instructions. The SNS chimaera was generated by fusing together N-terminal Sox19b (aa 1-53), Nanog HD (aa 195-254) and C-terminal of Sox19b (aa 113-284). The FNF chimaera was generated by fusing together N-terminal Fus (aa 1-299), Nanog HD (aa 195- 254) and C-terminal of Fus (aa 359-514). The NSN chimaera was generated by fusing together N-terminal Nanog (aa 1-194), Sox19b HMG domain (aa 195-262) and C-terminal Nanog (aa 263-392). A list of all primers used for cloning these constructs can be found below. All positive clones were validated by sequencing.

### List of all primers used for cloning

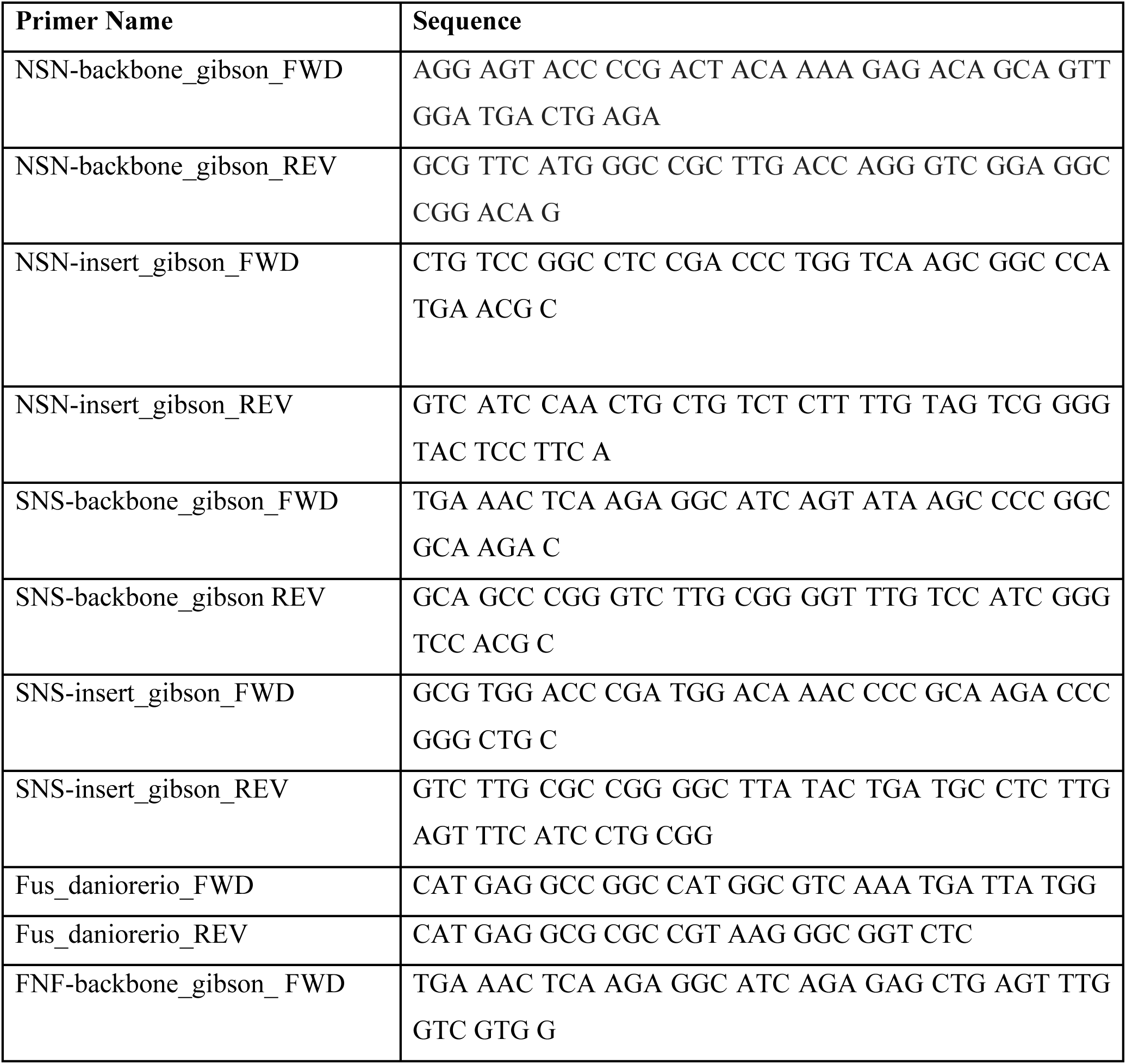
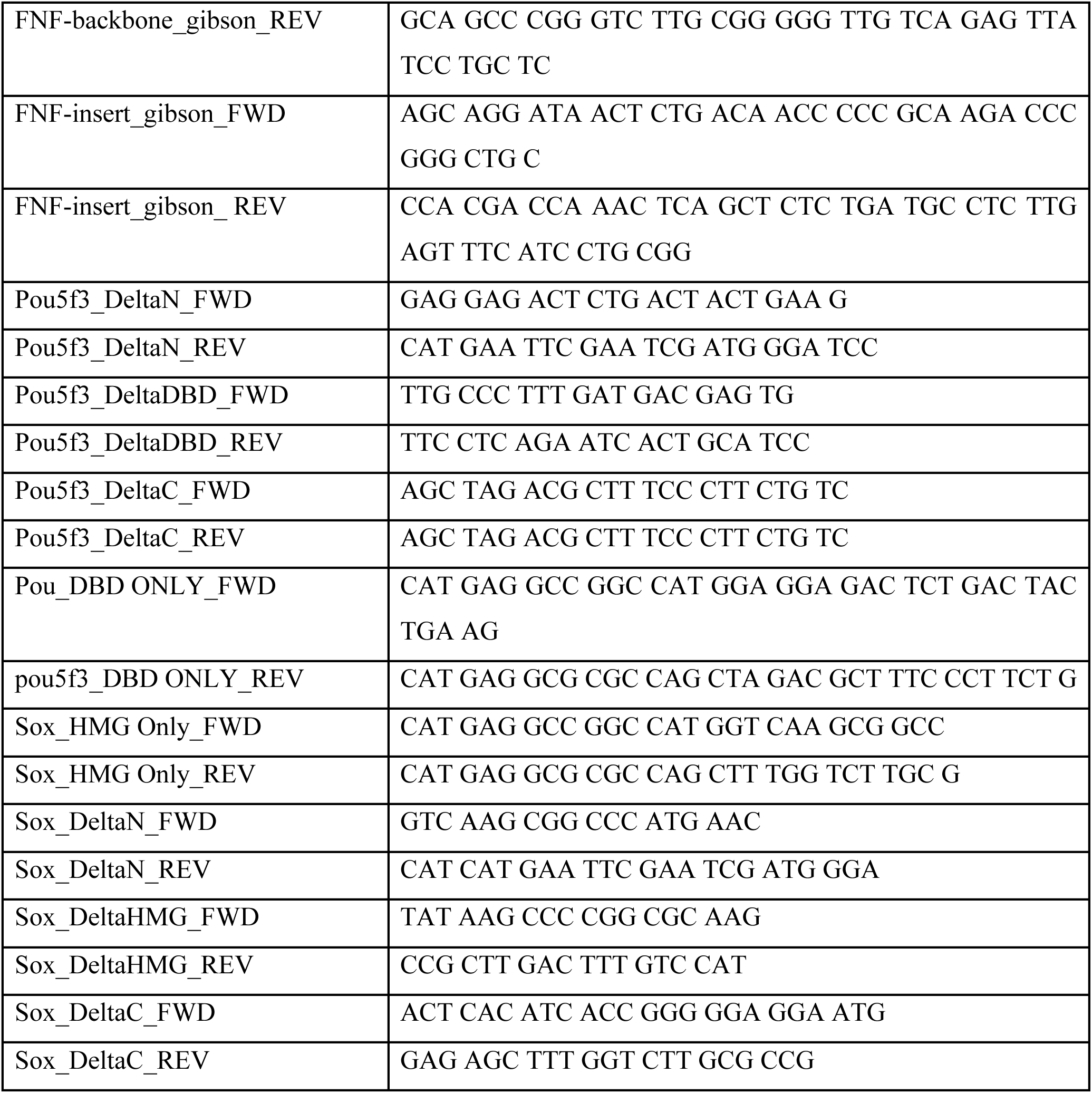

### mRNA production

Prior to *in vitro* transcription, 5-10 μg of plasmid was linearised with the restriction enzyme NotI (NEB, R3189L). The linearised plasmids were run on a gel to ensure complete digestion, and PCR purified using a PCR and gel purification kit, following manufacturer’s instructions (Qiagen, 28506). The SP6 mMessage mMachine *in vitro* transcription kit (Thermo Fisher Scientific, AM1340) was used for *in vitro* transcription, according to manufacturer’s instructions. 1μl of the mRNA was run on an agarose gel to check if a band of the correct size was obtained. The RNA was purified using Qiagen’s RNeasy MinElute kit, following manufacturer’s instructions (Qiagen, 74204). The purified product was diluted to a concentration of 600 ng/μl, aliquoted as 1μl aliquots and stored at −70℃ for several months.

### Genotyping mutant fish

Genotypes of maternal zygotic fish (*nanog^−/−^*, *sox19b^−/−^*, *pou5f3^−/−^*) were confirmed by phenotyping and/or genotyping. For genotyping, adult fish were finclipped and the genomic DNA was extracted according to the protocol described by Meeker and colleagues (35). Genotyping PCRs for the identification of Nanog, Sox19b and Pou5f3 mutants were performed as described previously (30, 36)

### Generating tetraploid embryos

Tetraploid embryos were generated using the HSII protocol (37) with some minor changes. Briefly, two waterbaths were maintained at 28℃ and 42.2℃. Beakers containing embryo medium (0.3x Danieaus) were placed in each water bath well in advance of embryo collection to allow the medium to equilibrate to the respective temperatures. Embryos were collected immediately after fertilization and transferred to a tea strainer, which was immersed in the embryo medium at 28℃. At 22 minutes post fertilization (mpf), the tea-strainer containing the embryos was transferred to the embryo medium at 42.2℃ for a heat shock for two minutes. Immediately after the heat shock, the embryos were transferred to a 28℃ incubator to recover. After 10 minutes of recovery, while the embryos were still at the 1-cell stage, they were microinjected as described above. Tetraploids were identified based on two parameters: (i) a delay of one cell cycle in the tetraploid embryos, when compared to embryos which were not subjected to heat shock (37), and (ii) counting the number of transcription bodies, labelled with MiR430 MOVIE(34).

### Live microscopy

#### Brightfield imaging

Bright-field images of whole embryos were taken with a stereomicroscope (Olympus, SZX-12) equipped with a CCD camera. The embryos were transferred to a glass dish containing embryo medium and manually dechorionated using forceps, prior to image acquisition. They were then positioned using forceps and a needle, taking care not to injure or pierce the embryos and imaged in a glass dish.

#### Mounting embryos for spinning disc microscopy

When embryos reached the 32-64 cell stage, they were transferred to a glass dish containing embryo medium and manually dechorionated using forceps, prior to mounting for imaging. Mounting medium was prepared by making a solution of 0.8% low melting agarose in Danieau’s, containing 15% v/v OptiPrep (Sigma Aldrich, D1156). The mounting medium was melted at 70℃, and then cooled to 37℃ in a glass vial, until the embryos were ready to be mounted. The dechorionated embryos were transferred to the mounting medium at 37℃ and mounted on an ibidi glass bottom μ-dish (Ibidi, 81158-400). The embryos were imaged using a Nikon CSU1 Yokogawa spinning disc microscope, in a temperature controlled chamber (Okolabs temperature controller and Nikon temperature controlled chamber), using a Nikon SR HP Plan Apo 100x / 1.35 Sil WD 0.3 objective. Serial optical sections were obtained for a z-stack of 30 μm, using intervals of 0.3 μm, with a time resolution of 3 minutes between sequential acquisitions.

### Image Processing and analysis

#### Software

All microscopy images were viewed and analysed using Fiji (Fiji is just ImageJ) (38). Cluster segmentation and analysis was performed using the Fiji plugin “3D Objects Counter”, except for the co-localisation analysis in Figure 2 (39). For Figure 2, the quantification of pairwise co-localisation of TFs was performed using Imaris. Figures were made using the Fiji plugin “ScientiFig” (40).

#### Cluster segmentation with Fiji

Individual nuclei at the indicated time points were segmented in 3D with a 250×250 pixel bounding box using Fiji. The resulting files were named with a nomenclature that allowed the nuclei to be traced back to the original files they were segmented from. TF clusters were segmented using the 3D objects counter plugin in Fiji (39) and manually curated. Clusters were segmented at the midpoint between two mitoses, except for Sox19b and its deletions and NanogΔN-mNG, for which clusters were segmented right after mitosis because these only from transiently. For full-length Sox19b-mScarlet and Sox19bΔN-mScarlet, which form only two clusters, clusters were counted manually, using maximum intensity projections of whole nuclei.

Co-localisation analysis of Nanog / Pou5f3 with Sox19b (Figure 2) was done in the early post-mitotic phase because Sox19b only forms clusters transiently. For the total number of Nanog / Pou5f3 clusters, we used the midpoint between two cell cycles when the maximum number of clusters was reached. We note that the Sox19b clusters always overlapped with the early appearing Nanog and Pou5f3 clusters.

#### Determining developmental stage in microscopy images

The embryonic developmental stages in microscopy images were determined on the bases of synchronicity of cell cycles, cell size, nuclear size, and inter-nuclei distance. The inter-nuclei distances provide a reliable method to determine cell sizes and can be found here (https://doi.org/10.5281/zenodo.8151889). The midpoint of each cell cycle was determined by measuring the time between two consecutive metaphases and taking the midpoint between these two timeframes.

### Data Analysis

#### Sample size

All experiments were performed keeping a minimum of three independent biological replicates. In these three biological replicates, multiple embryos—typically three or more per replicate—were included, consistent with standard practices in the field. The total number of embryos is represented by N. A minimum of 25 nuclei were selected for analysis (n>25). The nuclei were evenly distributed across embryos from all replicates to ensure balanced representation in the analysis. The details for the number of embryos (N) and number of nuclei (n) for each experiment are listed in the table below.

### Table containing information about the number of embryos (N) and number of nuclei (n)

| **Figure** | **Category** | **N (number of embryos)** | **n (number of nuclei)** |
| --- | --- | --- | --- |
| **1B** | Nanog diploid | 10 | 28 |
|  | Nanog tetraploid | 10 | 25 |
|  | Pou5f3 diploid | 9 | 30 |
|  | Pou5f3 tetraploid | 13 | 25 |
|  | Sox19b diploid | 12 | 38 |
|  | Sox19b tetraploid | 9 | 30 |
| **1D** | 2 pg | 8 | 25 |
|  | 10 pg | 9 | 25 |
|  | 20 pg | 11 | 45 |
|  | 40 pg | 14 | 53 |
| **S1** | Nanog 256-cell | 11 | 24 |
|  | Nanog 512-cell | 10 | 28 |
|  | Nanog 1k-cell | 9 | 23 |
|  | Pou5f3 256-cell | 9 | 24 |
|  | Pou5f3 512-cell | 9 | 30 |
|  | Pou5f3 1k-cell | 10 | 35 |
|  | Sox19b 256-cell | 6 | 12 |
|  | Sox19b 512-cell | 6 | 16 |
|  | Sox19b 1k-cell | 9 | 28 |
| **S2** | WT TLAB | 10 | 28 |
|  | *nanog*<sup>-/-</sup> | 8 | 27 |
| **S3** | 2 pg | 8 | 25 |
|  | 10 pg | 9 | 25 |
|  | 20 pg | 11 | 45 |
|  | 40 pg | 14 | 53 |
|  | 60 pg | 9 | 28 |
|  | 600 pg | 8 | 47 |
| **2B** | Nanog(-mNG) and Sox19b(-mSc) | 8 | 25 |
|  | Nanog(-mSc) and Pou5f3(-mNG) | 11 | 25 |
|  | Pou5f3(-mNG) and Sox19b(-mSc) | 10 | 25 |
| **3D** | Nanog FL | 8 | 27 |
|  | NanogΔHD | 9 | 30 |
|  | Nanog HD only | 8 | 30 |
|  | NanogΔN | 8 | 30 |
|  | NanogΔC | 10 | 31 |
|  | Pou5f3 FL | 9 | 39 |
|  | Pou5f3ΔDBD | 10 | 30 |
|  | Pou5f3 DBD Only | 8 | 30 |
|  | Pou5f3ΔN | 11 | 32 |
|  | Pou5f3ΔC | 7 | 39 |
|  | Sox19b FL | 9 | 31 |
|  | Sox19bΔHMG | 9 | 30 |
|  | Sox19b HMG Only | 10 | 30 |
|  | Sox19bΔN | 11 | 39 |
|  | Sox19bΔC | 9 | 30 |
| **3F** | Nanog FL | 10 | 28 |
|  | NanogΔHD | 7 | 30 |
|  | Nanog HD only | 9 | 30 |
|  | NanogΔN | 8 | 26 |
|  | NanogΔC | 8 | 40 |
|  | Pou5f3 FL | 9 | 30 |
|  | Pou5f3ΔDBD | 8 | 30 |
|  | Pou5f3 DBD Only | 9 | 30 |
|  | Pou5f3ΔN | 7 | 39 |
|  | Pou5f3ΔC | 9 | 32 |
|  | Sox19b FL | 11 | 38 |
|  | Sox19bΔHMG | 7 | 30 |
|  | Sox19b HMG Only | 9 | 30 |
|  | Sox19bΔN | 10 | 30 |
|  | Sox19bΔC | 8 | 30 |
| **4E** | Nanog FL | 9 | 28 |
|  | SNS chimaera | 11 | 40 |
|  | FNF chimaera | 9 | 25 |
| **4G** | Nanog FL | 3 | 180 |
|  | SNS chimaera | 3 | 275 |
|  | FNF chimaera | 3 | 389 |
|  | *nanog*<sup>-/-</sup> uninjected | 3 | 330 |

### Data Plotting and statistical analysis

The image analysis files in .csv format were exported to Rstudio software for data plotting and visualisation. The graphs were generated using the library ggplot2 (41). Statistical analysis was performed using GraphPad prism. Non-parametric statistical tests Mann-Whitney test (for 2 groups) or Kruskal Wallis test with Dunn’s comparisons (for multiple comparisons) were performed to calculate the p values for significance. Statistical significance was reported using a four-star system, with p < 0.05 indicated by (*), p < 0.01 by (**), p < 0.001 by (***), p < 0.0001 by (****), and non-significant results (p ≥ 0.05) denoted as ns.

## Data and materials availability

Raw imaging data are available upon request. All other data is available in the main text or the supplementary data.

## Acknowledgements

We thank members of the Vastenhouw lab for their support, helpful feedback, and stimulating discussions. We thank Noémie Chabot, Maria Cristina Gambetta, Arianna Penzo, Martino Ugolini, and Aleksandar Vještica for comments on the manuscript, and the following facilities and services for their support: UNIL – cellular imaging facility, fish facility, EPFL – BioImaging and Optics Core and fish facility.

## Author contributions

Conceptualization: S.D., N.L.V. Methodology: S.D. Investigation: S.D. Data analysis: S.D. Writing – original draft: S.D. and N.L.V. Writing – reviewing & editing: S.D. and N.L.V. Funding acquisition: N.L.V.

---

**Figure S1.**
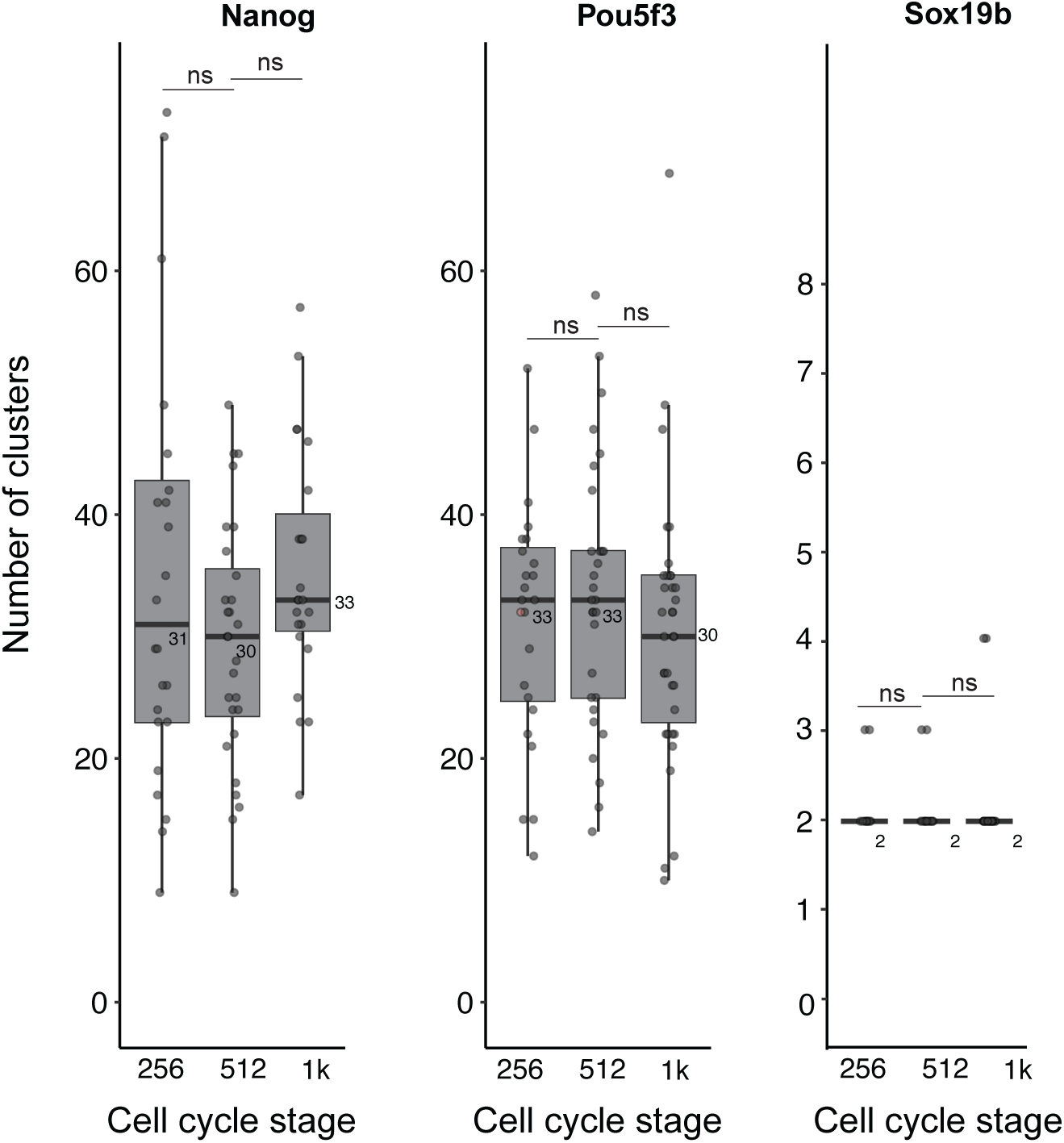
Quantification of TF clusters at 256, 512 and 1k-cell stage. Quantification of number of clusters for Nanog-mNG, Sox19b-mNG and Pou5f3-mNG at the 256-cell, 512-cell, and 1k-cell in WT TLAB embryos. The median values of the distributions are indicated in the graphs. With N as the number of embryos and n as the number of total nuclei, N ≥6 and n ≥12. Quantifications were performed at the midpoint between two mitoses, except for Sox19b for which the quantification was performed right after mitosis because clusters only from transiently. Statistical analysis was performed using Kruskal-Wallis test with Dunn’s multiple comparisons.

**Figure S2.**
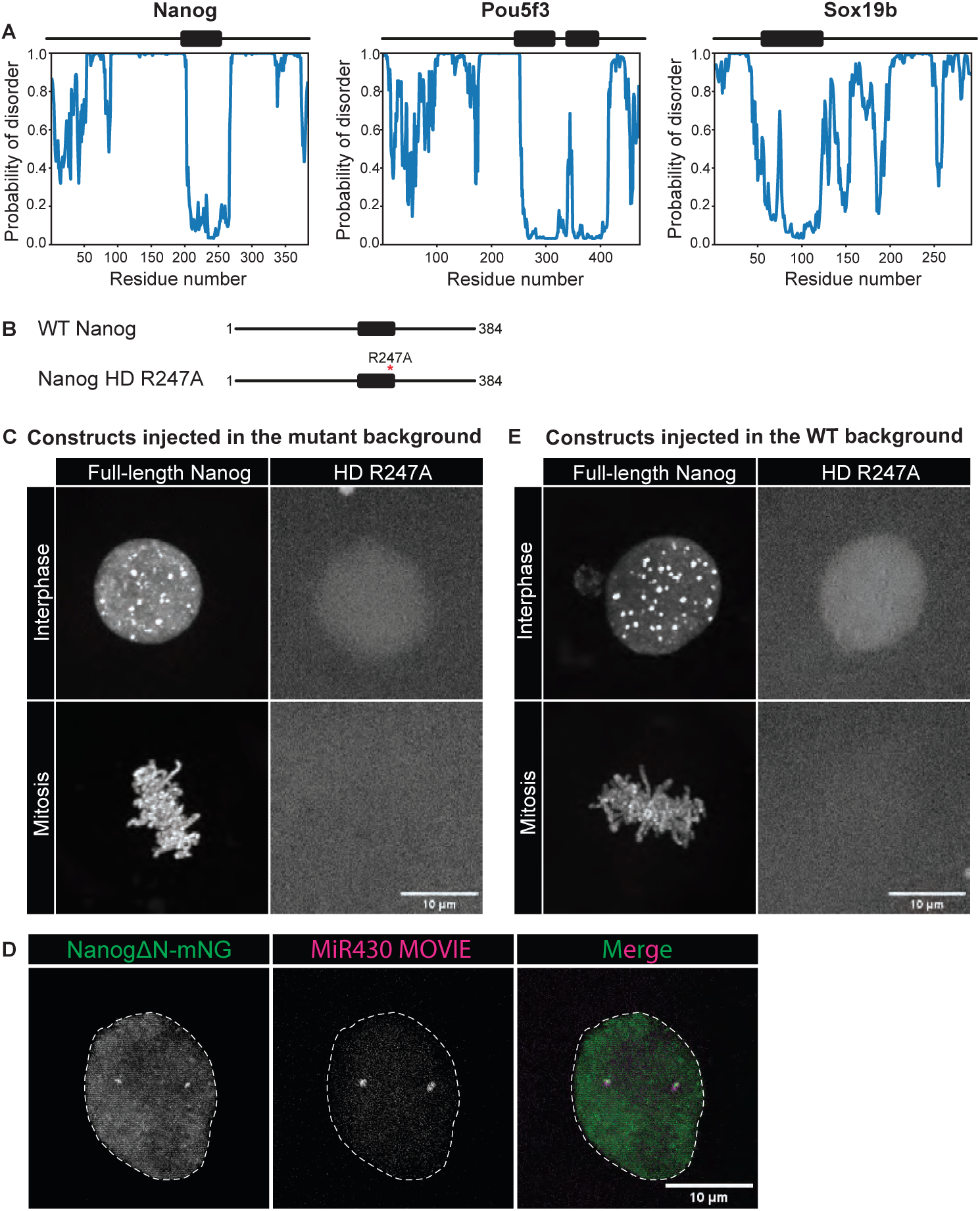
Additional information related to Figure 3. **A.** Disorder scan of Nanog, Pou5f3 and Sox19b generated using the ODINPred disorder prediction tool (1). Protein regions were classified as disordered if the disorder prediction score was above 0.5 and the region spanned more than 25 amino acids (aa). This scheme was used to generate the deletion constructs in Figure 3A. **B.** Schematic representation of full-length Nanog and the Nanog point mutant in which DNA binding is abrogated (Nanog HD R247A), for which RNA was injected in C and E of this figure. **C.** Images of MZ*nanog* embryos injected with WT Nanog and Nanog HD R247A. Shown are representative images of individual nuclei extracted from spinning disk confocal microscopy at 512-cell stage during interphase and mitosis. **D.** Colocalisation of NanogΔN-mNG and MiR430 transcripts in WT embryos. The images are taken right after mitosis because NanogΔN-mNG clusters only from transiently. **E.** Images of WT embryos injected with WT Nanog and Nanog HD R247A. With N as the number of embryos, and n as the number of nuclei. N ≥6 and n ≥20. In C-E, representative images of MIPs in Z of individual nuclei extracted from spinning disk confocal microscopy at 512-cell stage are shown.

**Figure S3.**
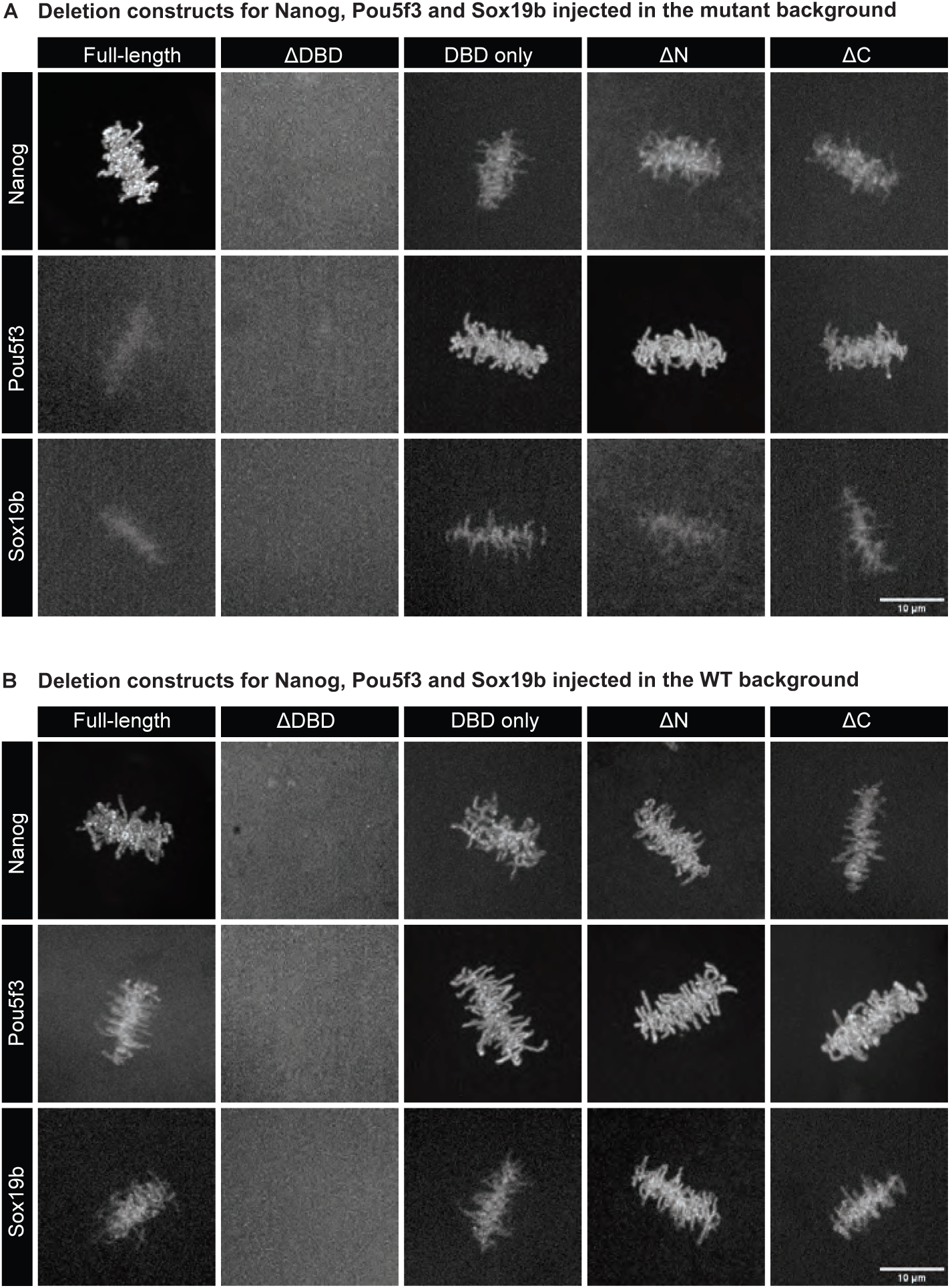
The binding of full-length and mutant Nanog, Pou5f3 and Sox19b proteins to DNA during mitosis. **A.** Images during mitosis of Nanog, Pou5f3 and Sox19B obtained after injection of the indicated constructs in the respective TF mutants. Related to Figure 3B, C. **B.** Same as in A but with constructs injected in a WT background, where endogenous protein of the injected factor is present. Related to Figure 3D, E. Shown are representative images of MIPs at mitosis extracted from spinning disk confocal microscopy.

**Figure S4.**
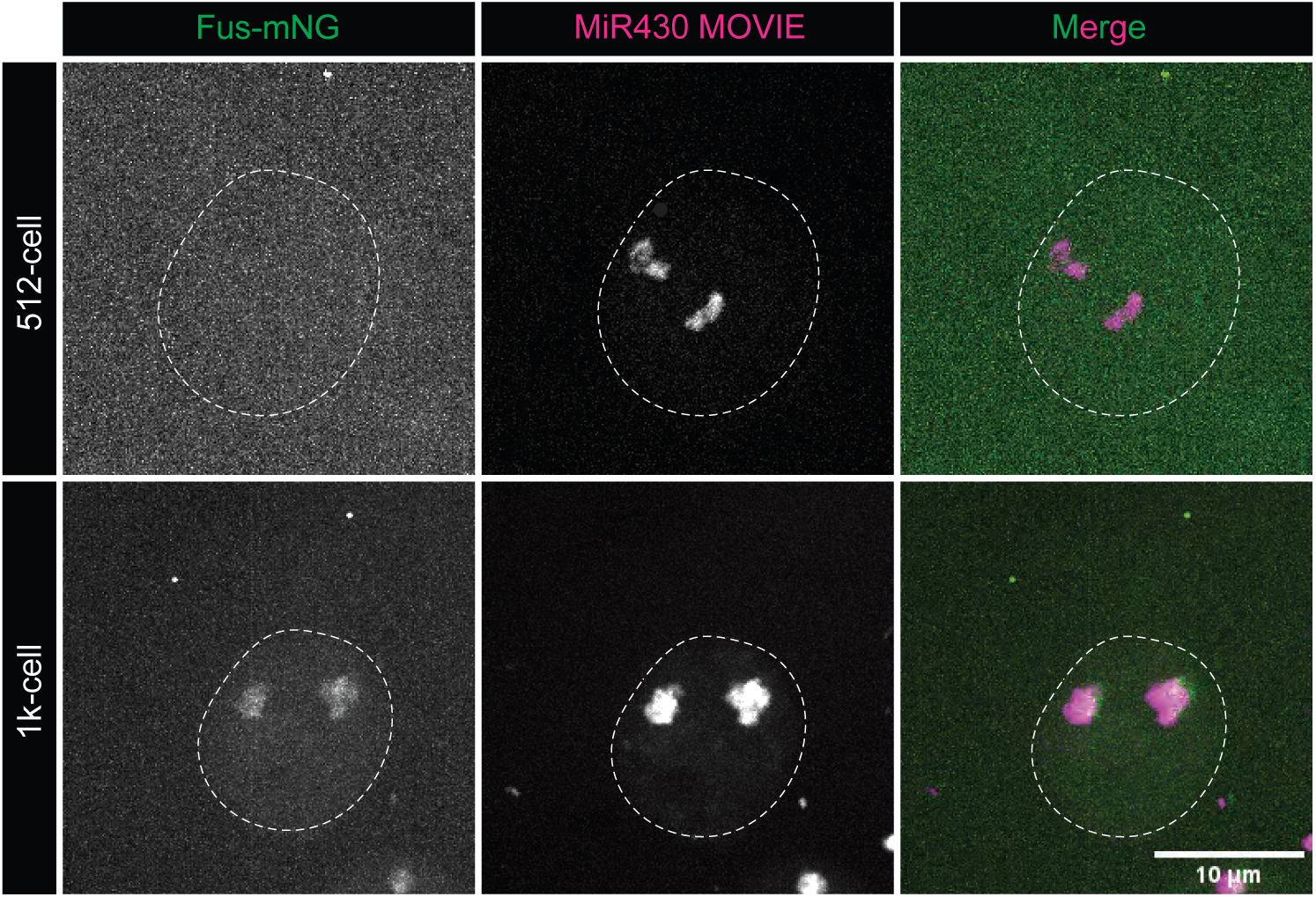
Fus-mNG does not form clusters at 512-cell stage. Visualisation of Fus-mNG and MiR430 transcripts in WT embryos at 512- and 1k-cell stage. At 512-cell stage, no Fus clusters can be detected. At 1k-cell stage, two Fus clusters can be detected. These colocalize with MiR430 transcripts. Shown are representative examples of individual nuclei extracted from spinning disk confocal microscopy during interphase. With N as the number of embryos, and n as the total number of nuclei N ≥5 and n ≥24.

**Figure S5.**
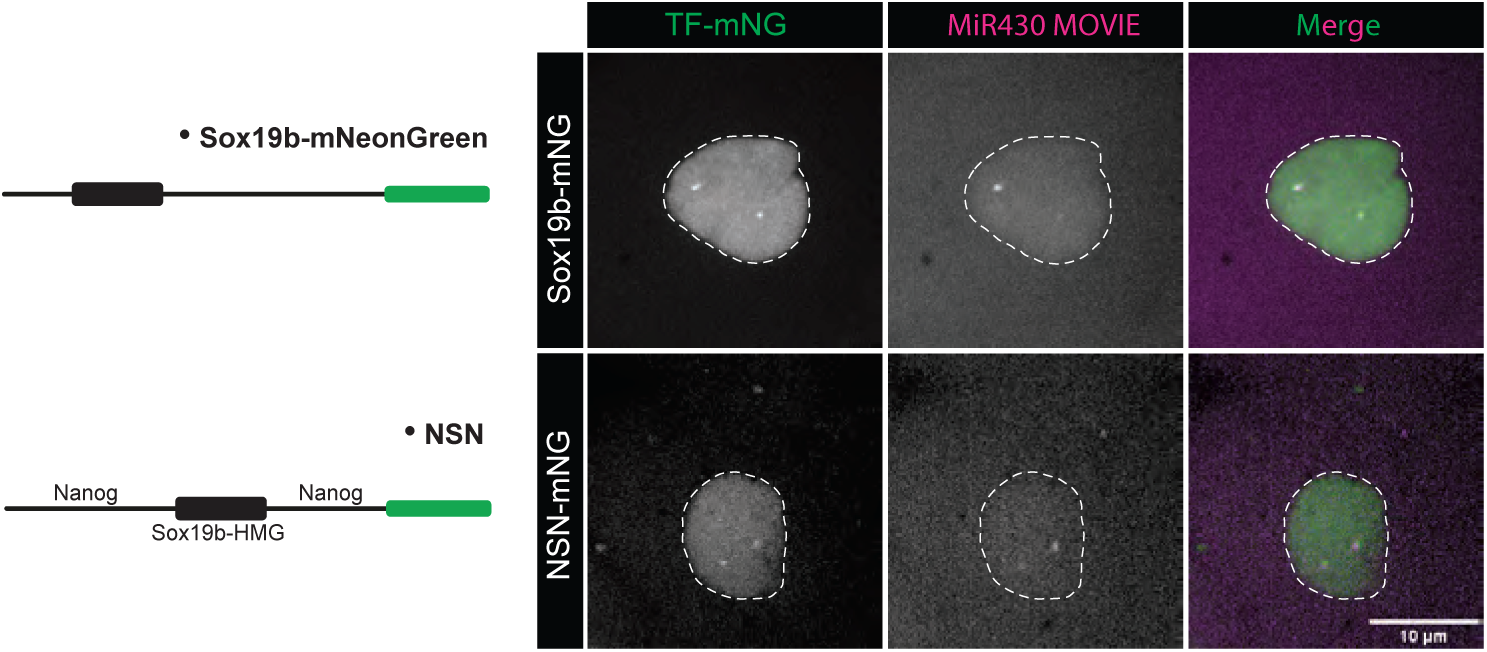
Specificity in clustering is mediated by the DBD (related to Figure 4). MZ*sox19b* mutant embryos were injected with FL Sox19b-mNG, or NSN-mNG, in both cases together with MiR430 MOVIE. Both proteins form two clusters and these colocalize with a marker for the *mir430* transcription bodies, MiR430 MOVIE. Shown are representative images of individual nuclei extracted from spinning disk confocal microscopy at 512-cell stage during interphase. Images were taken right after mitosis because these clusters only from transiently. With N as the number of embryos, and n as the number of nuclei N ≥6 and n ≥20.

